# Coding dynamics of the striatal networks during learning

**DOI:** 10.1101/2023.07.24.550305

**Authors:** Maxime Villet, Patricia Reynaud-Bouret, Julien Poitreau, Jacopo Baldi, Sophie Jaffard, Ashwin James, Alexandre Muzy, Francesca Sargolini, Ingrid Bethus

## Abstract

The rat dorsomedial (DMS) and dorsolateral striatum (DMS), equivalent to caudate nucleus and putamen in primates, are generally required for goal-directed and habit behaviour, respectively. However, it is still unclear whether and how this functional dychotomy emerges in the course of learning. In this study we investigated this issue by recording DMS and DLS single neuron activity in rats performing a continuous spatial alternation task, from the acquisition to optimized performance. We first applied a classical analytical approach to identify task-related activity based on the modifications of single neuron firing rate in relation to specific task events or maze trajectories. We then used an innovative approach based on Hawkes process to reconstruct a directed connectivity graph of simultaneously recorded neurons, that was used to decode animal behavior. This approach enabled us to better unravel the role of DMS and DLS neural networks across learning stages. We showed that DMS and DLS display different task-related activity throughout learning stages, and the proportion of coding neurons over time decreases in the DMS and increases in the DLS. Despite theses major differences, the decoding power of both networks increases during learning. These results suggest that DMS and DLS neural networks gradually reorganize in different ways in order to progressively increase their control over the behavioral performance.

## 1 Introduction

The dorsal striatum integrates sensory, motivational, and motor information transmitted by cortical and thalamic neurons, thereby enhancing the capacity to select actions that result in favourable outcomes, such as rewards, while also minimizing engagement in undesirable actions. Its anatomical projections from the cortex support a functional dissociation along the medio-lateral axis. The dorsolateral striatum (DLS) receives cortical inputs from the sensorimotor and premotor cortices, and was shown to sustain inflexible motor routines, stimulus-response learning and more generally habit behaviour [Balleine and O’Doherty, 2010, Balleine and Dickinson, 1998, Dickinson and Balleine, 1994, Yin et al., 2005, Dickinson et al., 1995, Graybiel, 1998, Packard and Knowlton, 2002, Yin et al., 2004, Hilario et al., 2012]. In contrast the dorsomedial striatum (DMS) is mainly targeted by associative and prefrontal cortices and supports goal directed behaviour, controlled by flexible action-outcome contingencies [Hunnicutt et al., 2016, Hintiryan et al., 2016, Hooks et al., 2018, Reig and Silberberg, 2014]. Accordingly, DMS and DLS display different task-related activity. The DLS was shown to develop preferential firing at the start and end of action sequences (‘task bracketing’) [Jog et al., 1999, Smith and Graybiel, 2013, Thorn et al., 2010, Barnes et al., 2005, Sales-Carbonell et al., 2018, Yin et al., 2009], whereas the DMS mainly activate at cue onset and movement onset, as well as with changes in reward prediction [Stalnaker et al., 2012] or changes in the reward delivery contingency [Regier et al., 2015]. This functional dichotomy between DMS and DLS is supposed to emerge in the course of learning, with the DMS being involved in the early stages and DLS’engagement becoming evident as animal performance approaches a plateau (Thorn 2010). Such gradual shift in striatal activity potentially supports a gradual change in learning strategy, from the execution of flexible and deliberative actions to optimized habitual motor sequences. But it remains unclear whether this gradual change is supported by a cooperation or a competition between the two striatal regions [Bicanski and Burgess, 2020]. Some studies have indeed demonstrated that the DMS and the DLS may work in competition [Malvaez et al., 2018], since DLS lesion facilitates the learning of DMS-dependent-goal-directed spatial strategies [Moussa et al., 2011], and DMS lesion accelerates the acquisition of inflexible motor sequences [Turner et al., 2022]. The aim of our study is to better characterize the interplay between DMS and DLS during learning processes, by monitoring the modifications of neural activity in the two structures at the level of single cells and neural networks, during the acquisition of a continuous spatial alternation task. In this task the rats learn to continuously alternate between the left and right arms of a T maze using the spatial cues available in the room. No specific cue provides any behavioral instruction, thus limiting experimentally-defined discrete actions that possibly influence ‘task bracketing’ activity. Moreover, this task requires the activity of both DMS and DLS from the beginning of learning, as we have shown in a previous study [Moussa et al., 2011]. It is therefore particularly suited for the study of DMS/DLS interplay during learning phases. DMS and DLS single neuron activity was recorded from the first day rats were exposed to the task until they reached and maintained a stable performance. We first monitored task-related activity across the different learning phases (from early acquisition to stable performance). Then, we decoded animal behaviour from DMS and DLS neural networks using classical analyses based on either firing rate or neural synchronization, but also combining the two measures using Hawkes process to reconstruct directed connectivity graphs of neurons. Altogether, these analytical approaches have provided a comprehensive description of the modifications of DMS and DLS neural networks across learning stages, thus allowing to unravel how their coding capacity modify in the course of learning. Overall, the results suggest that both DMS and DLS neural networks are engaged during all learning stages, in contrast with the common assumption of a gradual shift from DMS to DLS activity. Moreover, the results point to the importance of investigating neural network properties in addition to single neuron coding abilities, in order to better understand how brain structures participate to learning processes.

## 2 Results

### 2.1 Behavioral performance

Animal performance during each session was expressed as the number of paths traveled per minute. Paths consisted of 2 correct and 10 incorrect trajectories (see Methods Section and Figure 1C-D) and were defined based on a previous study ([James et al., 2023]). Performance evolution across training sessions was similar in all rats, with a gradual increase of the number of correct trajectories and a decrease of the number of incorrect trajectories (Figure 1B). Based on the rate of correct and incorrect paths (i.e. number of paths traveled per minute), we identified four successive stages of learning: stage 1, with a similar rate of correct and incorrect paths; stage 2, showing a progressive increase of the rate of correct paths and a progressive decrease of the rate of incorrect paths; stage 3, in which the rate of correct and incorrect paths stabilizes; and finally stage 4, showing almost exclusively correct paths.

**Figure 1:**
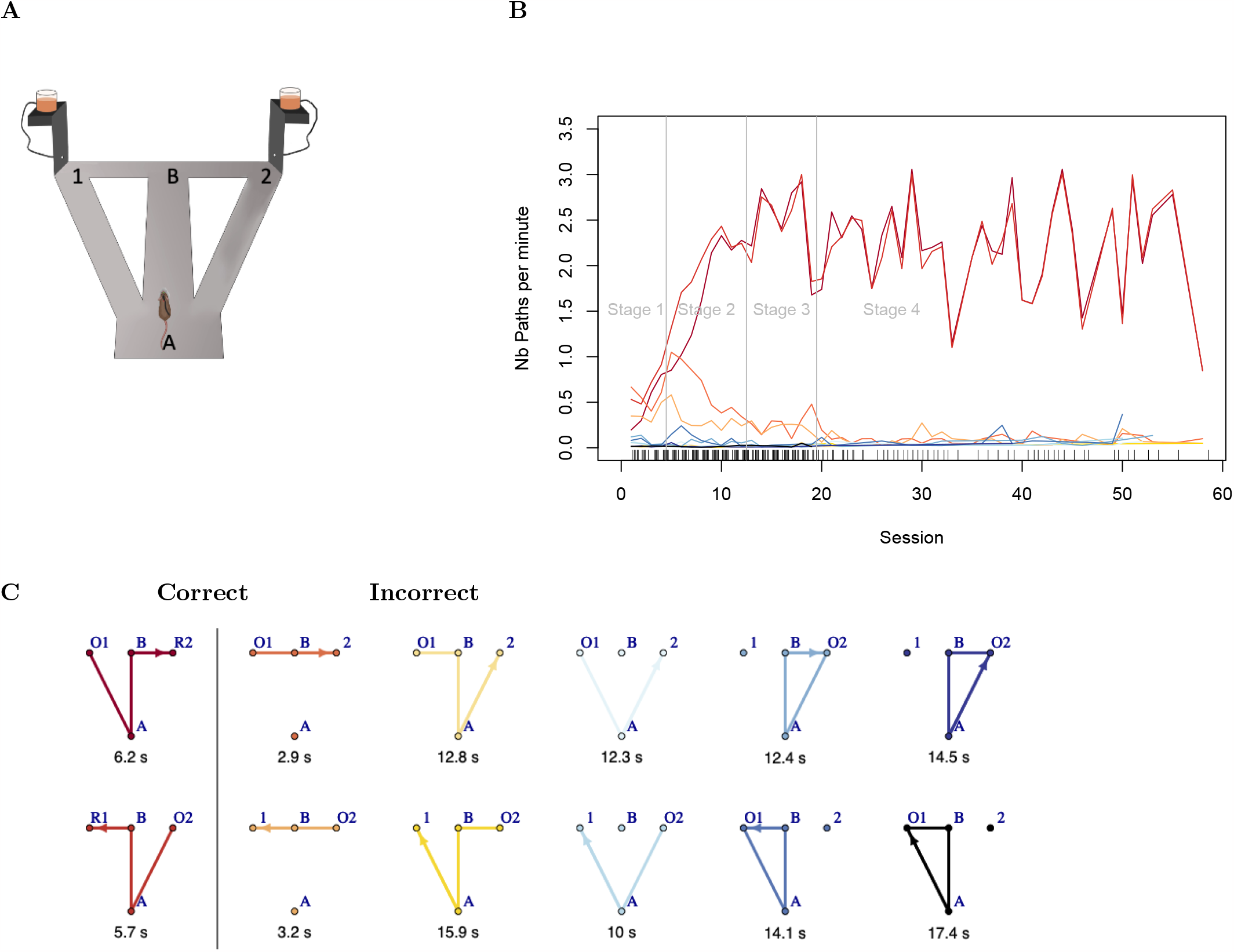
Behavioral data. **A**: Sketch of the maze with the two intersections A and B and the two feeders locations 1 and 2. **B**: Number of paths per minutes across learning stages. Each color corresponds to a path drawn with the same color in **C. C**: All the 12 considered paths with the 2 correct paths in the first column and the 10 incorrect paths in the second. Each path begins with an Onset O (1 or 2 depending on the feeder location). Only the correct paths end with a Reward R (1 or 2 depending on the feeder location). Under each path, the corresponding average duration in seconds for all rats and all sessions is shown.

### 2.2 Task-event coding neurons

We recorded 293 neurons in 4 DMS-implanted animals and 357 neurons in 3 DLS-implanted animals over a total of 206 training sessions (Supplementary Figure 6). On the basis of the firing rate and waveform properties (i.e., peak-valley distance and peak width at mid-height of the peak), cluster analysis revealed the presence of two clear clusters [Berke et al., 2004] (see Methods section and Supplementary Figure 8). Neurons with low firing rates and large waveforms were classified as putative medium spiny neurons (MSNs, GABA-ergic projection neurons), whereas the other group was identified as putative fast-spiking interneurons (FSIs, GABA-ergic interneurons). A total of 451 MSNs and 199 FSIs were found (Supplementary Figure 8). A chi-square test of independence showed no significant difference in the proportion of FSI/MSNs between DMS and DLS (p-val. = 0.97).

To analyze possible differences in firing activity in DMS and DLS neurons across learning stages, we first used a classical analytical approach and categorized neurons as coding task-events based on their firing activity in specific time windows. We identified six 500ms events: the two onsets O1 and O2 corresponding to the beginning of each path, the two intersections A and B of the maze and the two rewards R1 and R2, corresponding to the end of the correct paths (just before receiving the reward) (see Supplementary Figure 7 for a precise definition). Each of the six 500ms events was divided into five 100ms intervals. We classified as task-event coding each neuron whose firing activity in one 100ms interval varied significantly from all other intervals using a chi-squared detection method (see Methods Section for a description of the statistical identification procedure).

We found 65,5% of task-event coding neurons in DMS (16% non coding neurons) and 38% in DLS (24% non coding). 18.5% of DMS and 38% of DLS could not be classified because of too low firing rate. The Pearson’s Chi-squared test showed that the proportion of coding neurons is significantly higher in the DMS compared to the DLS (p-val.=7.57e-06). The results from the general linear model (with references: DLS and MSN) confirmed this effect (see the first column of Table 1). The proportion of task-event coding neurons in the DMS is greater than in the DLS (adj. p-val.=0.06). Interestingly, this proportion is stable across learning stages except for the stage 4 during which the proportion of DLS task-event coding neurons increases (adj. p-val.=0.07 with positive coefficient (coeff. = 0.68) at learning stage 4), while the proportion of DMS task-event coding neurons decreases (cross-effect adj. p-val. = 0.03, with negative coefficient (coeff = 0.57-1.86=−1.29)). Finally, in the DLS, MSN tend to code for task-events more than FSI (adj. p-val=0.03 with a negative coefficient (coeff. = −0.52)), whereas in the DMS the difference between FSI and MSN is marginal (cross-effect adj. p-val = 0.08 with coeff.=−0.52+0.57 = 0.05).

**Table 1:** Task-event coding neurons, neurons coding for left and right turns and neurons coding for full paths. First Line: On the top, number of coding neurons and below, number of non coding neurons, in the DMS (left) and DLS (right). The remaining lines report the results of a generalized linear model (see method section). The reference corresponds to DLS, MSN and learning stage 1. All p-values have been adjusted for multiplicity by Benjamini-Hochberg method. Only adjusted p-values ¡ 0.1 corresponding to a False Discovery Rate less than 10% have been reported with their associated coefficient.

| Coding type | Task-event | Left-Right Turns | Full Paths |
| --- | --- | --- | --- |
| $\frac{\# \text{Coding}}{\# \text{Non Coding}}$ DMS \| DLS | $\frac{192}{47}$ \| $\frac{136}{87}$ | $\frac{102}{144}$ \| $\frac{42}{172}$ | $\frac{226}{61}$ \| $\frac{182}{161}$ |
| DMS Coeff. (adj. pval) | 1.27 (0.06) | 1.12 (0.05) | 0.86 (0.03) |
| FSI Coeff. (adj. pval) | -0.52 (0.03) | - | - |
| Stage 2 Coeff. (adj. pval) | - | - | - |
| Stage 3 Coeff. (adj. pval) | - | 0.97 (0.06) | - |
| Stage 4 Coeff. (adj. pval) | 0.68 (0.07) | 0.98 (0.06) | - |
| Cross-Effect DMS:FSI Coeff. (adj. pval) | 0.57 (0.08) | - | - |
| Cross-Effect DMS:Stage 2 Coeff. (adj. pval) | - | - | - |
| Cross-Effect DMS:Stage 3 Coeff. (adj. pval) | - | -1.06 (0.08) | - |
| Cross-Effect DMS:Stage 4 Coeff. (adj. pval) | -1.86 (0.03) | - | -0.82 (0.08) |

Figure 2.A shows the normalized average firing activity and the corresponding confidence intervals at the different task-events, for neurons in the DMS (red) and DLS (blue) during the four learning stages. At the beginning of learning, during stage 1, the activity of DMS and DLS neurons decreases similarly during the R1 and R2 events. In stage 2, as behavioral performance improves, DMS and DLS neural activity starts to diverge. DMS neurons increase their firing rate particularly at the intersection B of the maze, before the rat turns into the left or right arm, whereas no significant change was observed at the intersection A, yet implying similar turning behavior. This result possibly indicates an involvement of the DMS in the implementation of the learning rule. As for the DLS, neurons change firing rate specifically at the reward areas during both R and O events. These modifications are similar to the DLS ‘task bracketing’ activity reported in previous studies [Thorn et al., 2010, Cunningham et al., 2021]. Finally, no specific task-related activity is observed in the DLS during the later stages of learning, whereas the DMS tends to show some activity at action boundaries similar to DLS, during stage 4, as also shown by Vandaele et al. [2019].

**Figure 2:**
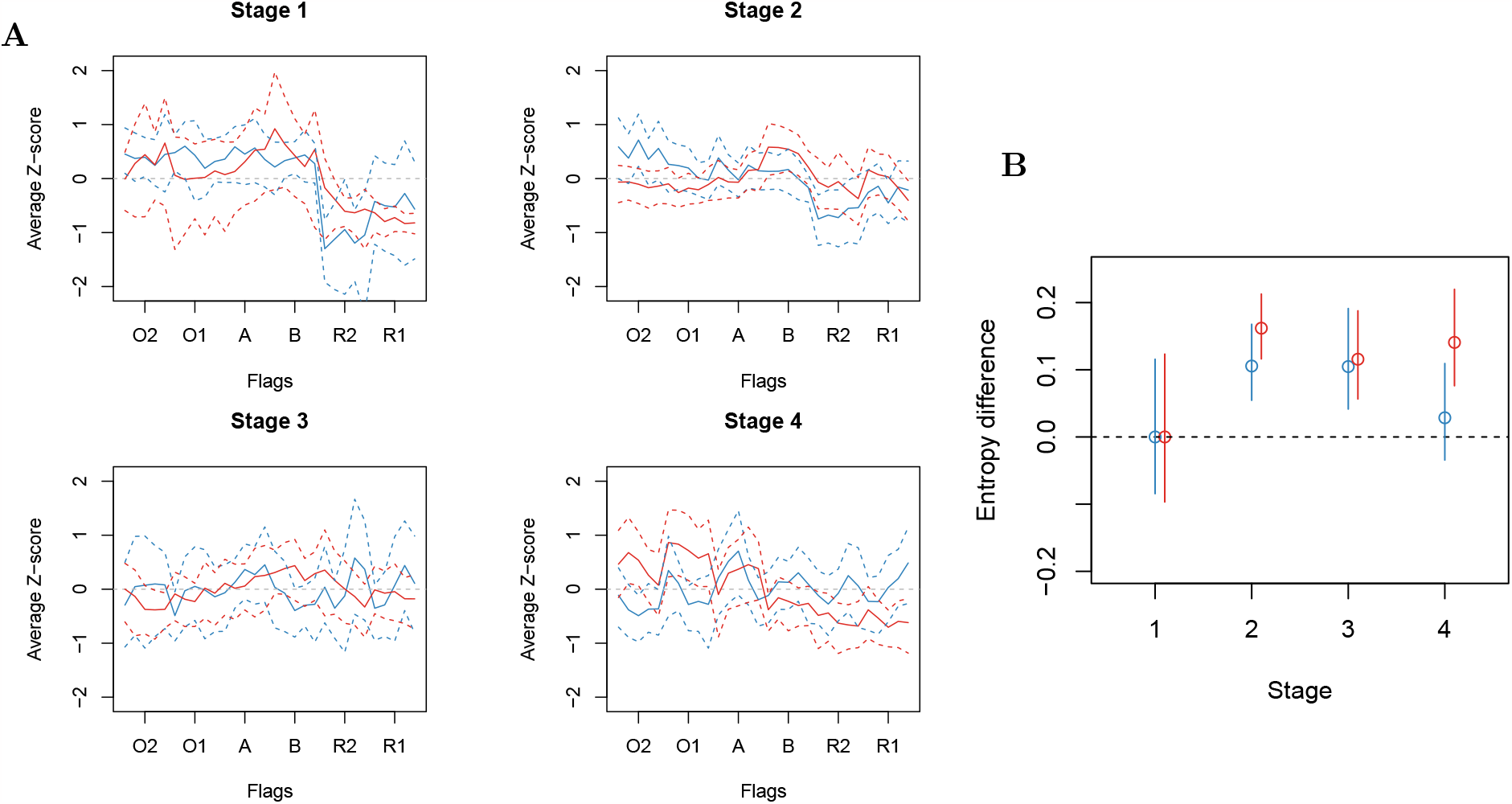
Z-score and Entropy. **A**: Average z-score of task-event coding neurons in DMS (red) and DLS (blue) in the four learning stages. Confidence bands at level 0.05 corrected for multiplicity over the 8 curves and the 30 time intervals (Bonferroni) are represented in dashed line. **B**: Difference in Z-score entropy between learning stages 2-to-4 and stage 1 (reference) for DMS and DLS. Confidence intervals at level 0.05 corrected for multiplicity over the 4 stages and 2 regions are pictured by a vertical line.

To test whether these changes in neural activity during task-events may reflect changes in coding capacity across learning stages, we estimated the z-score entropy from the firing rates of the task-event coding neurons in the DMS and DLS. A decrease in entropy reflects a decrease in the randomness of the (renormalized) firing rate in the neural population, which is classically interpreted as an increase in the coding ability of the neuronal population [Thorn et al., 2010]. As shown in Figure 2.B, the entropy of both DMS and DLS firing activity increases between stages 1 and 3, which may suggest a decrease in coding capacity during task acquisition for both areas. However, in contrast to the DMS, the entropy of the DLS tends to decrease in stage 4 to a level similar to that observed in stage 1. This may indicate that the DLS, but not the DMS, is involved in late learning stages following overtraining, as previously suggested [Tang et al., 2009]. However, given the overlapping confidence intervals, it is difficult to draw a clear conclusion. It should also be noted that in this task animals are free to perform any trajectory and no specific instructions are given at any time. Thus, an event-based analysis may not be the most appropriate to reveal learning-related changes in neural activities. We therefore analyzed DMS and DLS capacity to code for specific paths (rather than events) across learning stages, as presented in the next paragraph.

### 2.3 Neurons coding for left and right turns

In classical event-based analysis, a change in firing rate is considered as a code for a specific event in the task (i.e., movement onset, reward, choice), but not for the task itself (here for example dissociating left and right turns). For this reason, we compared the firing activity of each neuron in the central stem (from A to B intersections) between left-turn and right-turn trials, including correct and incorrect paths. If the firing activity is significantly different between left-turn and right-turn trials, the neuron is classified as a left/right turns coding neuron (see the Methods section for a full description of the statistical method). Table 1 (second column) summarizes the results of the statistical analysis. As for the task-event coding neurons, we detected significantly more left/right turns coding neurons in the DMS than in the DLS (adj. p-val= 0.05), but no difference between FSI and MSN. Moreover the proportion of coding neurons tends to increase during learning stages 3 and 4 in the DLS (coeff. 0.97 or 0.98 and adj. p-val=0.06), whereas it is less clear for the DMS (stage 3: cross effect adj. p-val. = 0.08, coeff =0.97-1.06=−0.09).

We then focus on neural synchronization as a marker of task encoding [Grammont and Riehle, 2003, Avital et al., 2013] and we calculated for each pair of DMS or DLS neurons recorded simultaneously in each session, the probability of being synchronized at intersections A and B. We analyzed a total of 1821 pairs of neurons. The cumulative distribution functions (c.d.f.) of all p-values (one per neuron pair) are plotted in Figure 3 for DMS (red) and DLS (blue) neurons, at each learning stage. If the neuron pairs are independent, the c.d.f. should be below the diagonal; any positive deviation from the diagonal, especially for small p-values, reflects a lack of independence (i.e., the neurons are synchronized). The Overall, these results demonstrate that DMS neurons code for left and right turns, particularly during initial learning, an activity that may support the acquisition of the task rule. This hypothesis is reinforced by the observation that DMS neurons are synchronized during initial learning, particularly before the rat chooses the left or right goal arm. In contrast, no specific synchronization pattern was observed in the DLS at any learning phase.

**Figure 3:**
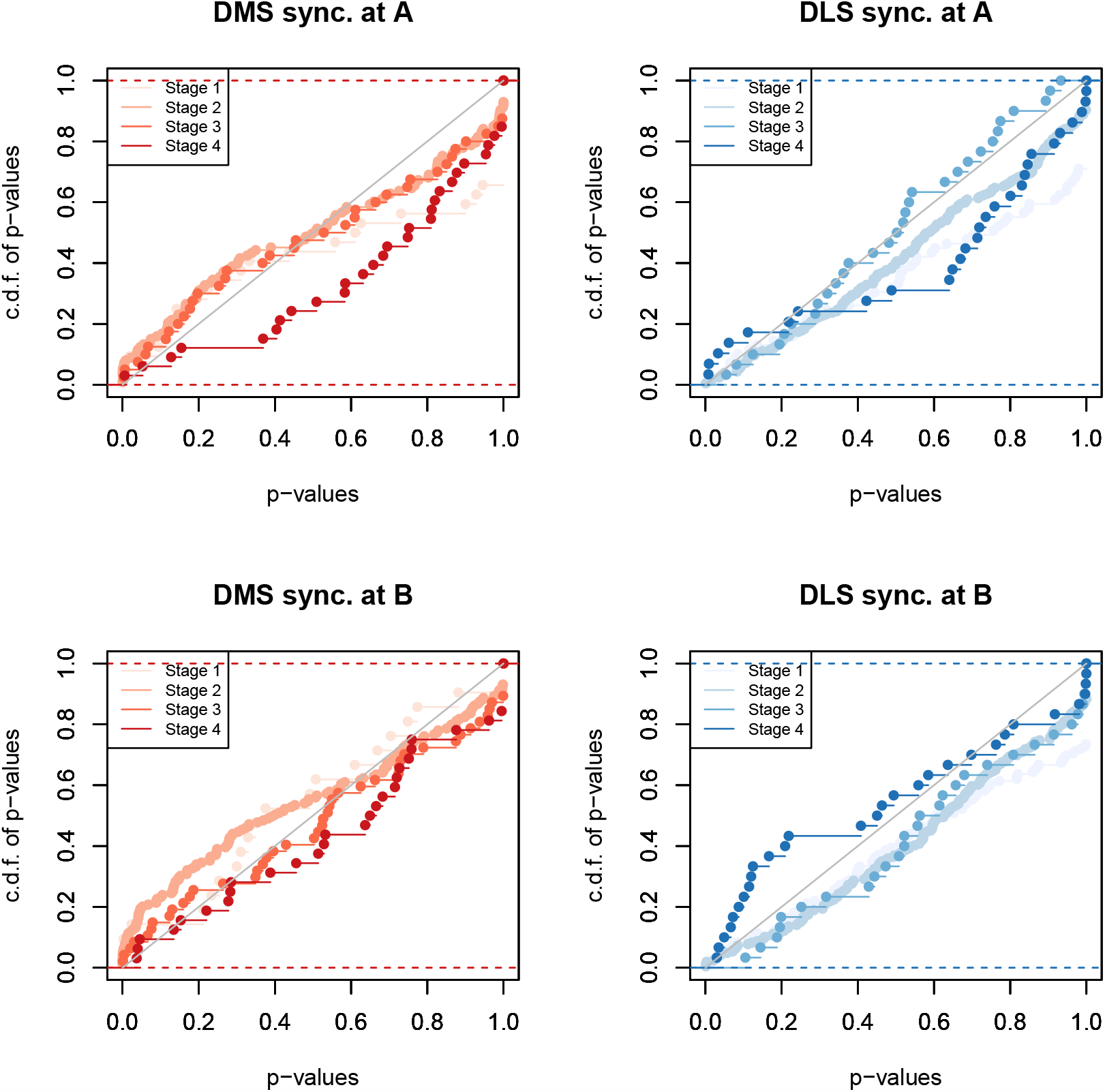
Synchronization analysis at intersections A and B. Synchronization between pairs of neurons was detected in 500ms time windows centered on the two maze intersections (A and B). Coincidences count was statistically compared to the empirical distribution of 50000 trial permutations. Cumulative distribution functions (c.d.f.) of the obtained p-values have been plotted as a function of the brain region (red for DMS and blue for DLS), intersection A or B and learning stages (color darkens as learning stage increases). A one sided Kolmogorov-Smirnov test of uniformity have been performed to compare all c.d.f. curves (16 in total). In addition, for each learning stage and maze intersection a two-sample one sided Kolmogorov-Smirnov test has been performed. All p-values were adjusted for multiplicity by Benjamini-Hochberg method. Note that c.d.f. of DMS p-values in stage 2 at intersection B is significantly above the diagonal (p.adj.= 8.610^−3^). Moreover, in stage 2 c.d.f. of DMS p-values at A and B are significantly above c.d.f. of DLS p-values (intersection A: adj. p-val. = 0.027; intersection B: adj. p-val. = 1.7 e-04) Kolmogorov-Smirnov test revealed a significant deviation from the diagonal for DMS neurons at stage 2, particularly at intersection B (p.adj = 8.6 e-03), whereas no significant effect was observed for DLS neurons at all stages. We then compared the c.d.f. of each learning stage between DMS and DLS to assess whether DMS neurons were more synchronized than DLS neurons. We found that in stage 2, the c.d.f. for DMS neurons was significantly higher than the c.d.f. for DLS neurons at both intersections A (adj. p-val. = 0.027) and B (adj. p-val.=1.71e-04). These results indicate that, overall, DMS neurons tend to fire simultaneously more than DLS neurons during initial learning. In addition, DMS neurons are significantly synchronized at intersection B of the maze, before the rat turns into the left or right goal arm, especially during initial learning.

### 2.4 Decoding animal behavior using Hawkes model

We then asked whether the previously identified coding properties were decisive for decoding animal behavior. In other words, we asked whether the information provided by neural activity was sufficient to guess the path taken by the rat (among the 12 identified). Neural activity is traditionally modeled by a Poisson process based on the firing rate. However, we have shown that the synchronization of neurons, in addition to the firing rate, also reflects their coding capacity: we therefore need a decoding model that takes into account both single neuron firing activity and neural interactions. We chose to use the multivariate Hawkes model, which includes neural firing rate, as well as the strength and directionality of the interactions between neurons[Lambert et al., 2018, Reynaud-Bouret et al., 2021]. Since our ultimate goal was to compare path decoding properties between DMS and DLS, we limited the analysis to the neurons showing significantly different firing activity between any two paths (181 DLS neurons and 226 DMS neurons) (see third column of Table 1). The proportion of path coding neurons in the DMS is again greater than in the DLS (adj.p-val. = 0.03), and tends to decrease over learning stages (cross-effect adj. p-val.=0.08 with coeff.=−0.82) similarly to task-event and left/right turns coding neurons (see Table 1). Since no significant difference was observed between MSN and FSI, both were included in the analysis.

Figure 4 shows the interaction graphs reconstructed from the Hawkes model versus the Poisson model (vertical line) during a representative session for a DMS-(red) and a DLS-implanted rat (blue). The reconstructed graphs for two different paths from the same session are shown on the same line. The color code represents either the firing rate for the Poisson model or the spontaneous part for the Hawkes model (i.e., baseline activity with all interaction functions canceled). For the Hawkes model, the directionality and strength of the interactions between two neurons are represented by the direction and color of the arrows, respectively. In this example, we can clearly notice that, for the same set of neurons, the graphs for the two paths are different while their firing activity is roughly the same.

**Figure 4:**
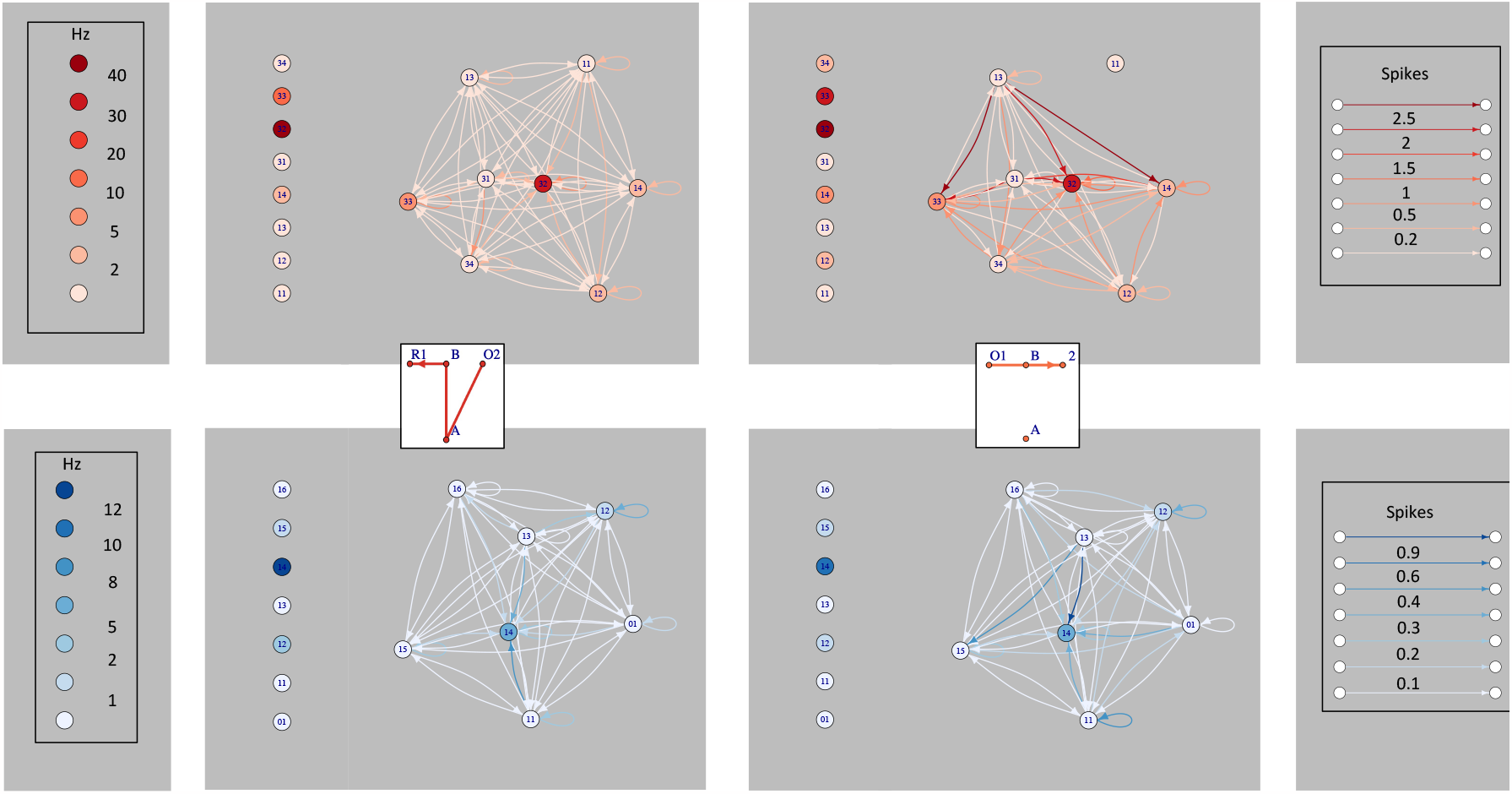
Examples of estimated network models for DMS (in red) and DLS (blue) neurons recorded from one DMS-implanted rat and one DLS-implanted rat during two training sessions (session 11 for DMS and session 6 for DLS) and two different paths (a correct path on the left and a straight path on the right). Each node represents one neuron. The color of the node represents either the average firing rate for the Poisson model (first vertical line on the left) and the spontaneous part for Hawkes model (with all interactions canceled). The color and directionality of the lines represent the interaction function. Right panel: color code for the interaction function, with the number of spikes modified by this interaction.

To evaluate the performance of the two models in decoding animal behavior, we first calculated the difference between the decoding power of the Poisson model, the Hawkes model, and a random guess. Decoding power is defined as the percentage of correct identification of the traveled paths, calculated by estimating the models on 2/3 of the paths in each training session and testing the model predictions on the remaining 1/3. As shown in Figure 5.A, the Hawkes model has better decoding power than the Poisson model, which predicts rat performance as well as a random guess. This may be due to the short path lengths (see Figure 1.B) combined with a fairly low firing rate of the neurons, which results in the prediction being based on 10-20 spikes per neuron in a given session. By taking into account other parameters of neuronal activity (i.e., the direction and strength of neuronal interactions), the Hawkes model is able to decode more efficiently the paths taken by the rats.

**Figure 5:**
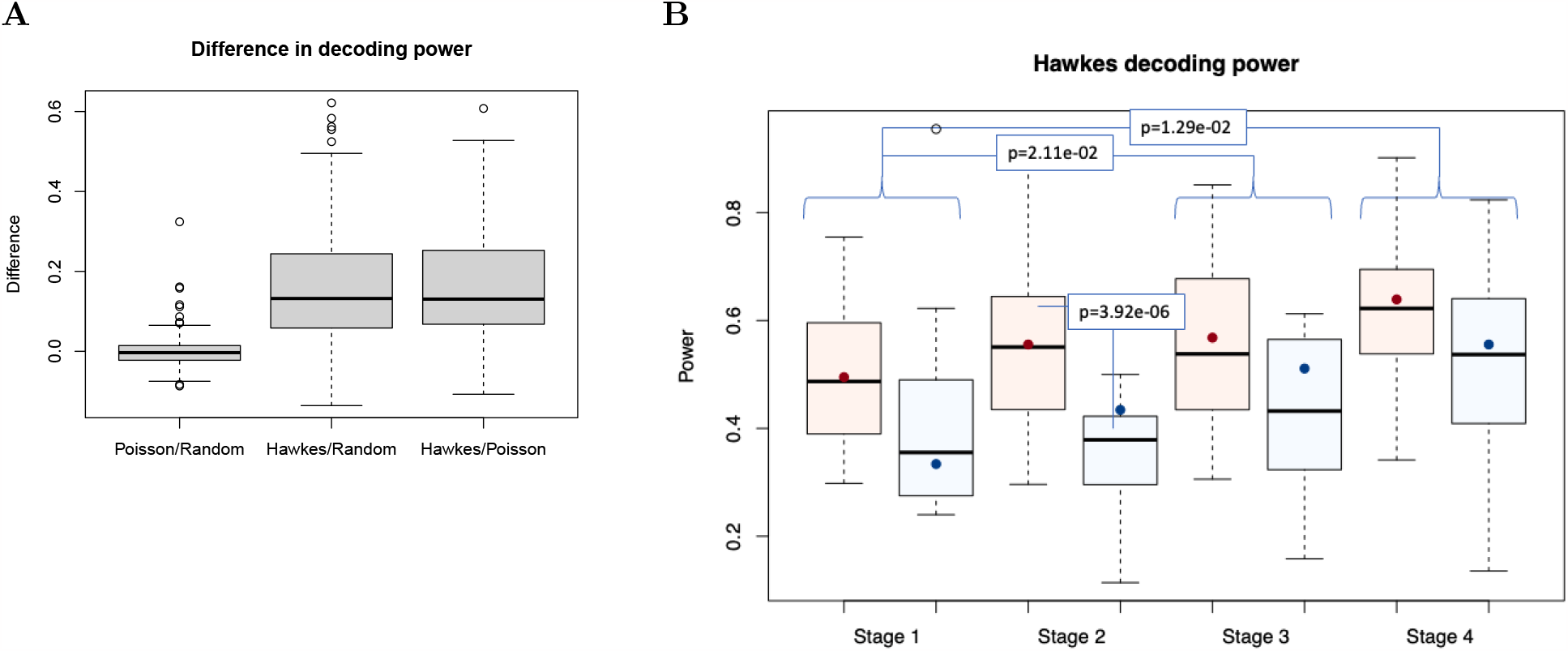
Decoding power. **A**: Boxplot showing the difference in decoding power (see Methods section for the precise definition) between the random guess (1/Nb of Paths), the Poisson model and the Hawkes model, for all rats and training sessions. **B**: Boxplot showing the Hawkes decoding power for DMS and DLS neurons across learning stages (red for DMS and blue for DLS). A beta regression with probit link is performed to compare DMS and DLS across learning stages. All p-values have been adjusted for multiplicity with Benjamini-Hochberg method. The colored dots represent the value predicted by the regression model for each boxplot. Significant adjusted p-values are indicated in the figure.

We then compared the decoding power computed by the Hawkes model in DMS and DLS across learning stages (Figure 5.B). To do so, we performed a beta regression with probit link (and explanatory variables: ‘number of paths’, ‘number of neurons’, ‘average firing rate’, ‘DMS-DLS’, ‘learning stage’ and ‘DMS-DLS’/’learning stage’ cross-effects) to disentangle the effects of interest (‘DMS-DLS’ and ‘learning stage’) from other auxiliary effects we need to consider. The results show that, in general, the decoding power in the later learning stages is greater compared to stage 1 (stage 3: adj. p-val.=2.11e-02; stage 4: adj. p-val.=1.29e-02). The number of paths (adj. p-val.=4.7e-12) and neurons (adj. p-val.=2.3e-07), as well as their average firing rate (p.adj= 1.3e-02) also have a significant effect, but no cross effect was observed. As for the comparison between DMS and DLS, the results show that the decoding power of DMS is globally greater than that of DLS (p.adj=2.8e-03), and particularly during learning stage 2 (adj. p-val.=3.92e-06). Note that the network size effect (number of neurons) has been taken into account in the model and cannot explain such difference. These results show that both the DMS and DLS neural networks undergo a gradual reorganization during learning, since the decoding power of the network increases as the rats performance improves in both DMS and DLS. However, it should be noted that DMS shows greater decoding power relative to the DLS during initial learning, possibly indicating that the acquisition of a spatially guided task rule without any temporal or cue instruction requires DMS neural activity.

## 3 Discussion

In this study we aimed at characterizing the dynamic interplay between DMS and DLS across learning phases, by analysing how neural activity at the level of single neurons and neural networks modifies in rats learning a continuous spatial alternation task. We showed that DMS and DLS display different task-related activity throughout learning stages. However, the progression of the coding ability across learning does not correlate with the capacity to decode animal performance. At the level of the network both structures increase their decoding power over time. But the DMS displays a greater decoding capacity in all learning stages compared to the DLS, despite a progressively decreasing number of task-coding neurons. Conversely, the smaller coding capacity of the DLS relative to the DMS is paralleled by an increasing number of coding neurons. These results suggest that the DMS and DLS neural networks differently reorganize during learning in order to progressively increase their control over the behavioral performance.

### DMS vs DLS single neuron coding properties across learning stages

We demonstrated that the DMS shows the greatest proportion of coding neurons across all learning stages, possibly indicating that it is more involved that the DLS in learning a spatial alternation strategy. During initial learning, DMS neurons tend to activate and synchronize at the intersection of the maze just before the rat choose to turn into the left or right arm. Neural synchronization has been related to stimulus encoding in different cortical areas [Grammont and Riehle, 2003, Avital et al., 2013] and can be possibly generalized to the encoding of abstract information such as the learning rule. Therefore, DMS activity at maze intersection possibly reflects its implication in the acquisition of the spatial rule, or could even sustains deliberative processes (i.e. decision making). It should be noted that previous studies have ascribed a similar role to the hippocampus and the prefrontal cortex as well as their interaction [Hassabis et al., 2007, Benchenane et al., 2010, Spiers and Gilbert, 2015]. Our result supports the hypothesis that DMS could be part of a network mediating decision making in spatially guided behaviors [Redish, 2016]. In contrast to the DMS, DLS neurons during early learning show mainly activation at action boundaries, as previously described in a cued (i.e. non spatial) T maze [Jog et al., 1999, Thorn et al., 2010] and in a lever-press task [Jin and Costa, 2010]. This ‘bracketing activity’ is supposed to improve the acquisition of behavioral routines and promote habit learning. Here we observe a similar ‘bracketing’ activity in the DLS but in the absence of any specific instruction, thus suggesting that a decomposition of the global task in multiple sub-tasks is at play even in the absence of any clear start-stop signals. This hypothesis is sustained by a recent study in which we showed that in the continuous T-maze task rats tend to use a behavioral strategy that consists in chunking sub-actions to perform complete paths [James et al., 2023]. Altogether these results point to a role of the DLS in identifying sub-actions or sub-tasks not exclusively in the context of habit learning.

As learning progresses, DMS and DLS single neurons show comparable coding properties, both activating at action boundaries in learning stage 4, similarly to what Vandaele et al. [2019] reported in a lever-press task. However, despite this progressive similarity, the activity of DMS and DLS neural populations strongly diverge across learning, since the proportion of coding neurons and the entropy of neural ensemble firing follow opposite trends in the two striatal areas. It is thus difficult to draw any clear conclusion based on single neuron firing properties. Therefore we decided to take advantage of a functional connectivity approach based on Hawked processes to infer neural network functions across learning.

### Decoding rat behavior from DMS/DLS network activities using Hawkes processes

One way to infer brain structure function in learning processes is to decode animal behavior from neural activity. Therefore, instead of looking for the conditions in which neural activity changes, we analysed how well we can reconstruct animal performance from neural activity. To do so, we used a classical Poisson model based exclusively on single neuron firing rate, and the Hawkes model that takes into account both single neuron firing rate and the interactions between neurons as well as the directionality of this interaction. This model allows to reconstruct a graph of functional connectivity between simultaneously recorded neurons, by analysing how much of the activity of a neuron is explained by the activity of the surrounding neurons. Therefore, both the strength of the connections between neurons and the direction of the interaction are represented in the graph. We observe that compared to the Poisson model the Hawkes model better decodes animal performance, indicating that neural interaction is a strong component of the striatal network function.

Using this analytical approach, we show that despite the progressive decrease in the proportion of coding neurons, the DMS network has higher decoding power than the DLS throughout learning stages. This result supports the hypothesis of a more selective implication of the DMS compared to the DLS in spatial alternation [Moussa et al., 2011]. However, it should be noted that the decoding power of both DMS and DLS increases over training, suggesting that as learning progresses neural activity in both areas conveys information necessary to support behavioral performance. It is interesting to note that the evolution of task-responsive neurons and decoding power across learning stages can be decorrelated. Specifically, the proportion of task-related coding neurons decreases in the DMS but the network itself exhibits enhanced decoding capacity. Conversely, the number of coding neurons and the decoding power increases over time in the DLS. Thus, an increasing proportion of coding neurons does not necessarily imply an increased ability to decode animal behavior from the same neural activities.Thanks to Hawkes processes, we are able to show that a neural network is capable of reorganizing itself to maintain or even increase its decoding power, and hence optimize behavioral performance, with a decreasing number of neurons.

In this context, it is necessary to define which behavior is decoded. Here we consider the decoded performance in terms of elementary paths, as described in Figure 1. In a recent study we have shown that rats learn a continuous alternation strategy by subdividing each paths into sub-paths or sub-actions [James et al., 2023]. Thus, the learning strategy consisted in chunking the different sub-actions, similarly to what was previously suggested for the acquisition of complex tasks [Botvinick et al., 2009]. We have tried to use the Hawkes model on the different sub-actions. However, given their short duration, the number of spikes per neuron was not sufficient to perform the analysis. In the future it would be interesting to design behavioral experiments where the sub-actions are long enough to allow to use the Hawkes decoding approach. Alternatively, it would be useful to increase the number of recorded neurons. In that regard, it should be noted that we obtained very good decoding results despite a relative low number of neurons recorded per session, which indicates that the striatal network is highly redundant [Reynaud-Bouret et al., 2021].

## Conclusions

In summary, we show that inferring neural network functions based exclusively on task-related single neuron firing activity is insufficient. By using the Hawkes model to decode behavior from neural network activities, we were able to achieve a more comprehensive description of DMS and DLS functions across learning. Altogether our results show that the coding pattern evolves differently in the two striatal areas over training, but both supports learning from early to late stages. .

It remains to be determined which is the exact nature of DMS and DLS interactions during learning. One possibility is that DMS and DLS activity supports distinct learning strategies and control action selection through a competitive mechanism. The results we obtained at early stages of learning supports this hypothesis, but the increased engagement of the two areas across learning stages is not coherent. Alternatively, we may speculate that either DMS- and DLS-mediated action selection process occurs exclusively at the beginning of the learning, or that action selection is operated by the upstream cortical areas, and the differential striatal properties testify (and possibly amplify) such process. Once the behavioral strategy has been implemented, both striatal areas may be implicated in maintain the performance, that may explain why the decoding power in both areas increases over training.

## 4 Methods and Materials

### 4.1 Animals and surgery

Seven male Long-Evans rats (Janvier, Le Genest-St-Isles, France) weighing 300-350 g were housed in individual cages (40 cm long x 26 cm wide x 16 cm high) with food and water ad libitum and maintained in a temperature-controlled room (20°C +/- 2). One week after their arrival, animals were handled daily by the experimenter for 7 days. 4 animals were then implanted with tetrodes aimed at either the left (3 rats) or the right (1 rat) dorsomedial striatum (DMS) at the following coordinates: AP: *±* 1mm, ML: ±2.2 mm from the midline, DV: −3mm below the dura. Two animals were implanted with tetrodes aimed at the left dorsolateral striatum (DLS) and one rat was implanted bilaterally at the following coordinates: AP: *±*1mm, ML: ±3.7 mm from the midline, DV: −3mm below the dura 6). The surgery was performed under sterile conditions and under general anaesthesia (Ketamine 75 mg/kg (Imalgene 1000, Merial, France)/Medetomidine 0.25 mg/kg (Domitor, Janssen, France)). As postoperative treatment, the rats were injected with antibiotic (Clamoxyl, 150 mg/kg) and analgesic (Tolfedine, 4 mg/kg). After surgery, the rats were given 5-7 days of recovery. They were then subjected to a food deprivation program that kept them at 90 % of their body weight during behavioral testing. All experiments were performed in accordance with the National Institute of Health’s Guide for Care and Use of Laboratory Animals (NIH Publication no. 80-23) revised in 1996 for the UK Animals (Scientific Procedures) Act of 1986 and associated guidelines or the Policy on Ethics approved by the Society for Neuroscience in November 1989 and amended in November 1993 and under veterinary and National Ethical Committee supervision (French Agriculture Ministry Authorization)

### 4.2 Microdrives and recording setup

Four tetrodes formed a bundle threaded through a piece of stainless-steel tubing. Each tetrode consisted of four twisted 25μm nichrome wires. The connector, tubing and wires could be moved downwards by turning the drive screw assemblies cemented to the skull. Cable was connected to the rat’s headstage, which contained a field effect transistor amplifier for each wire. The signals from each tetrode wire were amplified 10 000 times, bandpass filtered between 0.3 and 6 kHz with Neuralynx amplifiers (Neuralynx, Bozeman, MT, USA), digitised (32 kHz) and stored by the DataWave Sciworks acquisition system (DataWave Technologies, Longmont, CO, USA). A red light-emitting diode (LED) attached to the head assembly was used to determine the position of the rats. The LED was filmed by a CCD camera mounted on the ceiling above the maze, and its position was tracked at 25 Hz by a digital point tracker.

### 4.3 T maze apparatus

The T-maze consisted of four 10 cm wide, grey-painted wooden tracks (with walls of 2 cm height on each side), a 100 cm long central stem, a 100 cm long crossbar forming the two goal arms, and two additional tracks each connecting the distal end of one goal arm to the base of the central stem. The reward wells were located at the distal end of each choice arm. Food rewards (45 mg sugar pellets) were released from two feeders (MedAssociates) mounted above the wells and activated by remote manual switches. The maze was elevated 40 cm off the ground on a metal frame. The apparatus was illuminated by four symmetrical spotlights (40 W) mounted on the ceiling. A centered radio above the maze was used to mask uncontrolled disturbing sounds and the experimenter was located in an adjacent room. Behavioral training was performed as follows. After one week of recovery period from surgery, the rats were familiarized with the maze in daily 20-minutes sessions for two days, during which they were allowed to freely explore the apparatus and collect randomly scattered sugar pellets. Training began on the third day with either one or two 20-minutes sessions per day. Animals were gently placed at the intersection point labelled ‘A’ in Figure 1.A of the ‘continuous T-maze’ and let free to explore each arm as they wished. In order to obtain a 15 mg sugar pellet, they had to run along the central stem and alternately enter the left or right arm of choice. Rats were submitted to one training sessions per day. Tetrodes were lowered by either 25 *μ*m or 50 *μ*m after each training session or every two sessions, for a maximal length of 2mm (between 1 and 2mm, Supplementary Figure 6).

### 4.4 Data Analysis – behavior

Animal position (x/y coordinates) over time was sampled at 25 Hz. A preprocessing of the data, consisting mainly in clearing rapid head movements and reflection artifacts of the camera, was first performed. We identified 12 possible elementary paths (i.e. 2 correct paths and 10 incorrect paths) that rats could travel along the maze (Figure 1.C). Note that some rare portions of the trajectories were not classified as elementary paths and were removed from the analysis. These mainly include animals jumping out of the maze or jumping in a non contiguous spot in the maze. Animal performance was expressed as the number of paths (either correct or incorrect) per minute travelled by each rat during the training sessions. Based on rat performance, we identified four successive learning stages : Stage 1, with a similar number of correct and incorrect paths; Stage 2, showing a progressive increase of the number of correct paths and a progressive decrease of the incorrect ones; Stage 3, in which the number of correct and incorrect paths stabilize; and finally Stage 4, showing almost exclusively correct paths (Figure 1.C). The duration of each path type was also scored.

### 4.5 Data analysis – neuronal activity

Neural data from the four DMS implanted rats were also used in another study investigating rat learning strategies using credit assignment models [James et al., 2023]. All the results reported in the present study are original.

#### Spike Sorting

Spike sorting was performed manually using the graphical cluster-cutting software Offline Sorter (Plexon). Units selected for analysis had to be well discriminated clusters with spiking activity clearly dissociated from background noise. Units that were lost or whose waveform changed too much before the session was completed, were excluded. Units having interspike intervals ¡2 ms were removed due to poor isolation, as were cells with a peak firing rate ¡1 Hz. A total of 650 cell clusters was accepted (293 DMS and 357 DLS units). Since the tetrodes were lowered at the end of the recording session, neural activity in each session should be collected from different neurons. However, if two neurons recorded in two successive sessions showed similar characteristics in their firing patterns, waveforms and clusters, they were included only once.

#### MSN and FSI classification

For each neuron we computed the log of the global firing rate, the peak-valley distance (PV) (i.e. the distance between the peak and next minimal value) and the peak width at half-height (W) from the digitalized waveforms. These three variables were next centered and renormalized. Then we used the hierarchical clustering method with ‘Ward 2’ distance to build the dendrogram, showing clearly the existence of 2 clusters (Supplementary Figure 8). The k-means cluster algorithm with *k* = 2 was used to identify the clusters. The cluster with small firing rate and large PV and W corresponded to putative medium-spiny neurons (MSN), whereas points with large firing rate and small PV and W were allocated to the putative fast-spiking interneurons (FSI) cluster.

#### Identification of coding neurons

To identify coding neurons we performed three different analysis that were based on either temporal or spatial firing activity. In the first analysis, a task-event coding neuron was defined as a neuron showing significantly different firing activity during any of the six task-related temporal events (described below). For the second analysis (left/right coding neurons), we focused on the neural firing within the central stem of the maze with respect to whether the rat will subsequently chose the left or right goal arm. In this case, a coding neuron should activate differentially for left and right turns. The third analysis allowed us to categorize path coding neurons as neurons showing significantly different firing activity between any two of the 12 paths described in the behavioral section. Path coding neurons were then used in the decoding analysis. The results from these analysis were used to compare the proportion of coding neurons across learning stages, brain areas (DMS - DLS) and types of neurons (FSI - MSN). Whatever the type of coding, the detection of coding neuron is the same and is described hereafter.

#### Chi-squared method for detection of coding neurons

The purpose of this statistical analysis is to test whether the firing activity is constant over all possible conditions (null hypothesis) or if it is significantly different in at least one given condition (i.e. a 100ms bin or a particular path) with respect to all other possible conditions. These possible conditions will include the 30 temporal bins defining the six task events in the first analysis (i.e. five 100ms-bins per event), or the two left and right paths for the second analysis, or the paths taken in a session for the third analysis. Given *N*_*k*_ the number of spikes produced in a given condition *k* = *k*, …, *K*, we compute the p-value of the chi-square test that decides if (*N*_1_, …, *N*_*K*_) is a multinomial distribution of parameters 

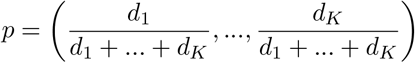

and 

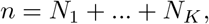

with *d*_*k*_, the duration of the condition *k* in the session. The test gives a valid p-value if *np*_*k*_ *≥* 5 for all coordinates *k*. Therefore if it is not the case, we can either drop a condition or group together conditions. If despite that, there is still not enough spikes, the corresponding neuron was removed from the analysis.

All neurons with a Benjamini-Hochberg (BH) adjusted p-value lower than 0.05 were declared as coding for a particular set of conditions. To assess the difference in the proportion of coding neurons (see Table 1) we then used a generalized linear model (Bernoulli with probit link) to explain the coding character as a function of brain region, neuron type and learning stages (with cross effects Brain region-Learning stage and Brain region-neuron type). Each time, DLS, MSN and Stage 1 were used as references. In a preliminary study, firing rates and all other possible cross-effects were added to the model, but since no significant results even without correction were found, they have been discarded. The fact that the firing rate does not impact the coding character is in adequation with the chi-square detection method: as soon as there are enough spikes in total in the session to use the method, there is no bias *per se* toward largest firing rates, once the FSI character has been taken into account.

- *Task-event coding neurons.* To identify event-based coding properties, we analyzed neural firing during specific time windows in four areas of the maze : the two intersections (depicted as A and B in Figure 1.A) and the two reward areas (depicted as R1 and R2 in Figure 1.A). We defined six 500ms time events based on the passages through the different areas (see Supplementary Figure 7). For the two intersections A and B, we extracted the central 500 ms time window of the trajectory within the two areas. For the reward areas we defined two different events: the reward obtention (R1 and R2 for the left and right goal arm) including the first 500ms after the rat entered the reward areas, and the movement onset (O1 and O2 for the left and right goal arm) including the last 500ms before the rat left the reward areas (Supplementary Figure 7). For each neuron we calculated the average firing rate during the 500ms events by refining the time window in 5 bins of 100ms. A task-event coding neuron is a neuron that has been detected by the chi-square method to show a firing rate activity in one 100ms bin that is different over all 30 time bins (i. e. 5 bins for each of the 6 events). If the condition *np ≥* 5 is not fulfilled, we have grouped the 100ms bins into 6 conditions, corresponding to 6 time windows of 500 ms around each event (intersections, rewards, onsets). If the condition is still unfulfilled the neuron is discarded. The z-score of an event coding neuron *n* in a 100ms bin of time *t* is given by 

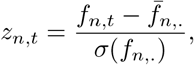

where *f*_*n,t*_ is the firing rate of neuron *n* in bin *t*, 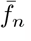,. is the average over the bins and σ(*f*_*n*,._) the corresponding standard deviation over the bins. Then, we averaged the normalized (z-score) activity of DMS and DLS event coding neurons over each neuronal population - brain region^*^ stage - (see Figure 2.A). We also estimated the entropy of the spike probability distribution in each population from the ensemble z-scores across learning stages by counting the number **Nz**_*I*_ of *z*_*n,t*_ in a given population that appears in an interval *I* of z-score given by segmenting [*−* 3.5, 5.5] in bins of 0.5 length, then the entropy of the distribution of z-score was estimated by 

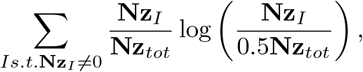

where 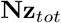 is the total number of z-score in the population. Note that 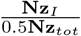 estimates the density of the z-score distribution inside interval *I*.
- *Neurons coding for left and right turns.* Here we focused on DMS and DLS neural firing activity in the central stem of the maze (between A and B intersections, Figure 1.A) with respect to whether the rat will subsequently choose the left or right goal arm. Here the chi-square method is performed only with two conditions and there is no grouping or dropping of conditions if *np ≥* 5 is not fulfilled. In this case, the neuron is just discarded.
- *Neurons coding for paths.* Firing activity in DMS and DLS during any of the twelve possible paths was calculated. We use again the chi-square method to identify significant spike activity in one of the twelve paths over all paths. If any of the path *k* does not satisfy *np*_*k*_ *≥* 5, this condition is simply left out. This is in particular the case if *d*_*k*_ = 0, meaning that this path is not taken by the rat during the session. If even with only two paths left, the condition *np ≥* 5 is not satisfied, the neuron is discarded.

#### Neuronal synchronization at maze intersections

For each learning session, we performed a trial shuffling / permutation analysis to detect synchronization of spiking activity between pairs of simultaneously recorded neurons at the intersection areas of the maze A and B (see Albert et al. [2015, 2016]). More precisely, for any pair of neurons simultaneously recorded, we isolated the number of times the rat passed through A or B and for each passage we extracted the central 500 ms window, defined as a trial. The number of coincident spiking activity observed within 20ms in each trial was compared to the empirical distribution obtained by computing 50 000 permutations of the trials, to obtain a p-value. We then calculated the cumulative distribution functions (c.d.f.) of all p-values from pairs of neurons in the DMS and the DLS across learning stages. We quantified the deviation of the c.d.f. from the diagonal by a Kolmogorov-Smirnov test of uniformity and the comparison between DMS and DLS c.d.f.s by a two-sample Kolmogorov-Smirnov test.

#### Decoding behavior using Poisson and Hawkes models

Decoding of rat behavior (i.e. choosing a particular path among the 12 identified, Figure 1.C) was implemented using a Poisson and Hawkes model. We chose Poisson model since neural firing is generally modeled in that way. In addition, we used the less conventional multivariate Hawkes model because it includes not only firing rate but also the strength and the directionality of the interaction between pairs of neurons [Lambert et al., 2018, Reynaud-Bouret et al., 2021]. Since we focused on path decoding, we included in the analysis exclusively DMS and DLS path coding neurons. In particular, this prevents the method to be diluted by a different proportion of non coding neurons in each area.

The Poisson model is defined by an intensity (instantaneous firing rate) of neuron *n* in condition (path) *k* given by 

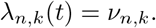

The parameter *v*_*n,k*_ represents the firing rate of neuron *n* in condition *k*. Hence if *N* is the total number of path coding neurons in the session, the Poisson model has *N* parameters per condition.

The Hawkes model is defined by an intensity of neuron *n* in condition (path) *k* given by 

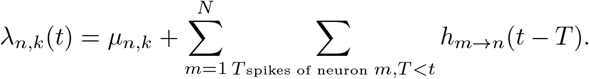

The parameter *μ*_*n,k*_ represents the spontaneous firing rate of neuron *n* in condition *k*. In addition, the interaction *h*_*m*→*n*_ quantifies the impact of one spike of neuron *m* at time *T* on the apparition of a spike on neuron *n*. The functions *h*_*m*→*n*_ are decomposed into 6 plateaux of length 10 ms each associated with a different value *a*_*m,n*,1_, …, *a*_*m,n*,6_, so that the value of *h*_*m*→*n*_ between 10 and 20 ms is for instance *a*_*m,n*,2_. The strength of an interaction is then measured by 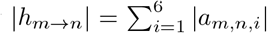. In total, *N* + 6*N* ^2^ parameters per condition are necessary for a Hawkes process. In particular, it includes self-interactions.

In each model, the parameters are estimated by minimizing the least-square contrast that is 

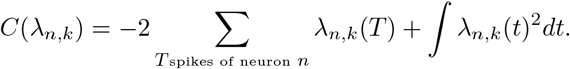

More precisely, for Figure 4, the sum and integral above are considered over all time *T* or *t* such that condition *k* holds. The estimated parameters *v*_*n,k*_ for the Poisson and *μ*_*n,k*_ for the Hawkes are given by the color of the nodes. The strength of the interaction *h*_*m*→*n*_ is given by the color of the edge *m→ n*.

The computation of the decoding power is more intricate. In each session, we consider only the paths that are taken at least 3 times. We defined as a trial each single path (and the corresponding neural activity) taken by the rat. So for condition (path) we have at least 3 trials. For each condition *k*, we computed estimators of the two models by using only 2/3 of the trials, leading to estimators of the intensity 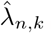. To do so, it is sufficient to restrict the integral and summation in the least-square contrast *C*(*λ*_*n,k*_) = *C*_2/3_(*λ*_*n,k*_) to time inside 2/3 of the trials. For each condition *k*, it remains 1/3 of the trials. For each remaining trial, we compute the least-square contrast 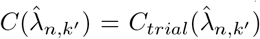, for all condition *k*′, where now the time in the sum and integrals in *C* is restricted to this precise trial. Therefore 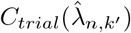 measures the similarity between the model estimated in the condition k’ and the spike train of the trial under inspection, which is in condition k. The estimated condition of a given trial is therefore given by 

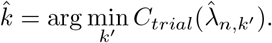

The decoding power of a model is the proportion of good guess, that is the proportion of trials for which 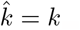.

The decoding power strongly depends on the number of paths (Nb Paths) within a session. Indeed, if there is not enough information in one path to decode rat behavior, the decoding power should be equivalent to 1/ Nb Paths. Therefore, we used this value as reference (i.e. random guess), that was used to assess the differences in decoding power between Poisson and Hawkes model in Figure 5.

Then, we calculated the decoding power for each striatal region across the four learning stages. As previously noted, the decoding power depends on the number of paths (Nb Paths) but also on the number of (path coding) neurons (Nb Neurons) and their average firing rates (Firing). Moreover, the encoding power is a quantity in [0, 1] that is best modeled by a beta variable. Therefore we performed a beta regression with probit link to test the effect of Nb Paths, Nb Neurons, Firing, Striatal Region (i.e. DMS/DLS), Learning Stage and cross effects Region/Stage, to disentangle the effects of interest (Region, Stage) and the other auxiliary effects that we need to take into account.

## Acknowledgments

This work was supported by the French government, through the UCA^Jedi^ and 3IA Côte d’Azur Investissements d’Avenir managed by the National Research Agency (ANR-15-IDEX-01 and ANR-19-P3IA-0002), by the interdisciplinary Institute for Modeling in Neuroscience and Cognition (NeuroMod) of the UniversitéCôte d’Azur and directly by the National Research Agency ANR-08-JCJC-0125-01 and ANR-19-CE40-0024 with the ChaMaNe project. It is part of the [Computabrain](https://neuromod.univ-cotedazur.eu/computabrain) project.

## Supplementary Figures

**Figure 6:**
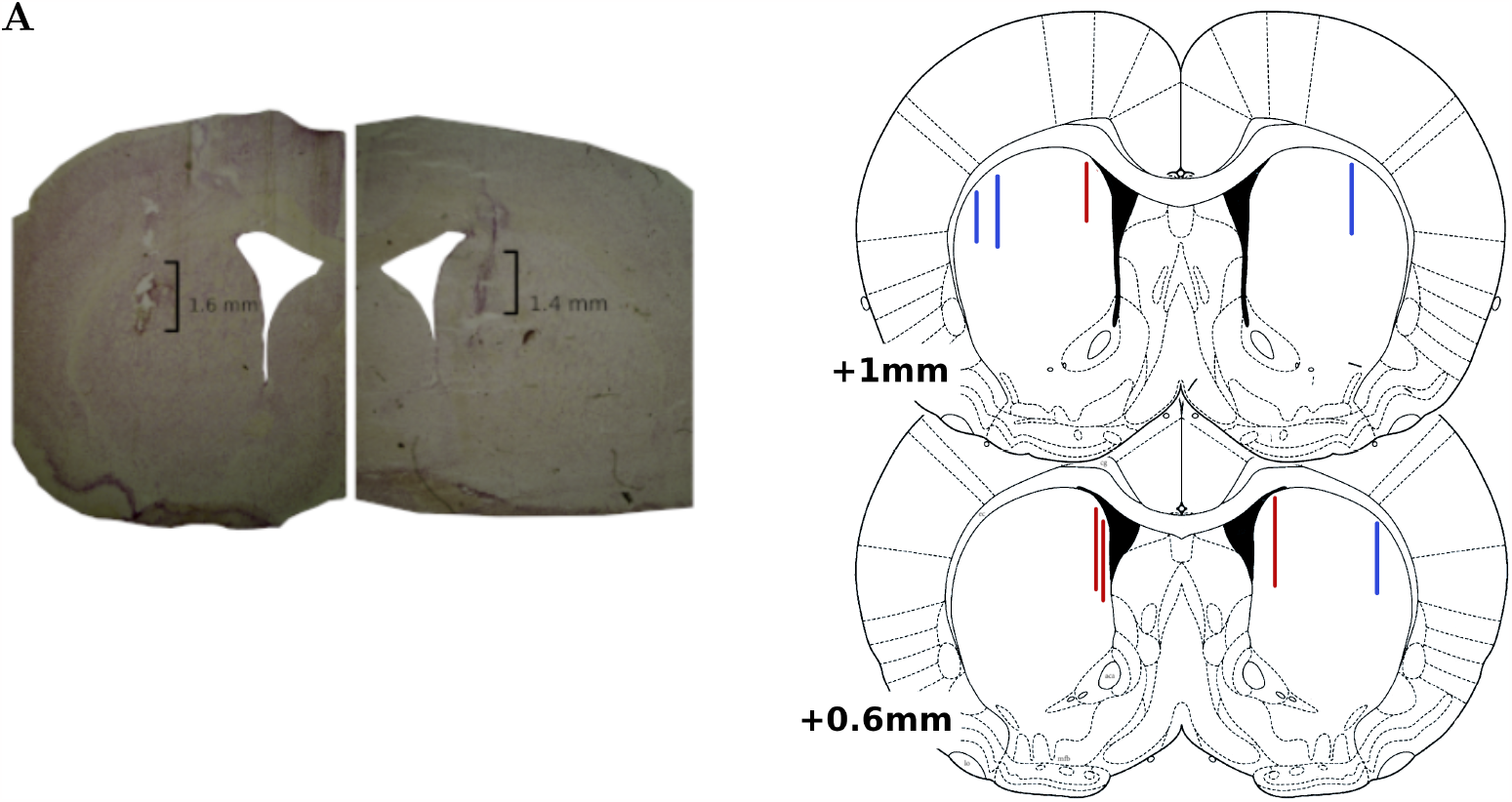
Reproduction of tetrode positions and recording sites in the DMS and DLS. **A**: Photos of Nisl-stained coronal sections from two representative rats implanted in the DLS (left panel) and DMS (right panel). The tetrode tracks are visible in both sections. The vertical black bars represent the lengths of striatal tissu from which neural recordings were performed. **B**: Representation of the recording sites from all rats (DLS in blue and DMS in red)

**Figure 7:**
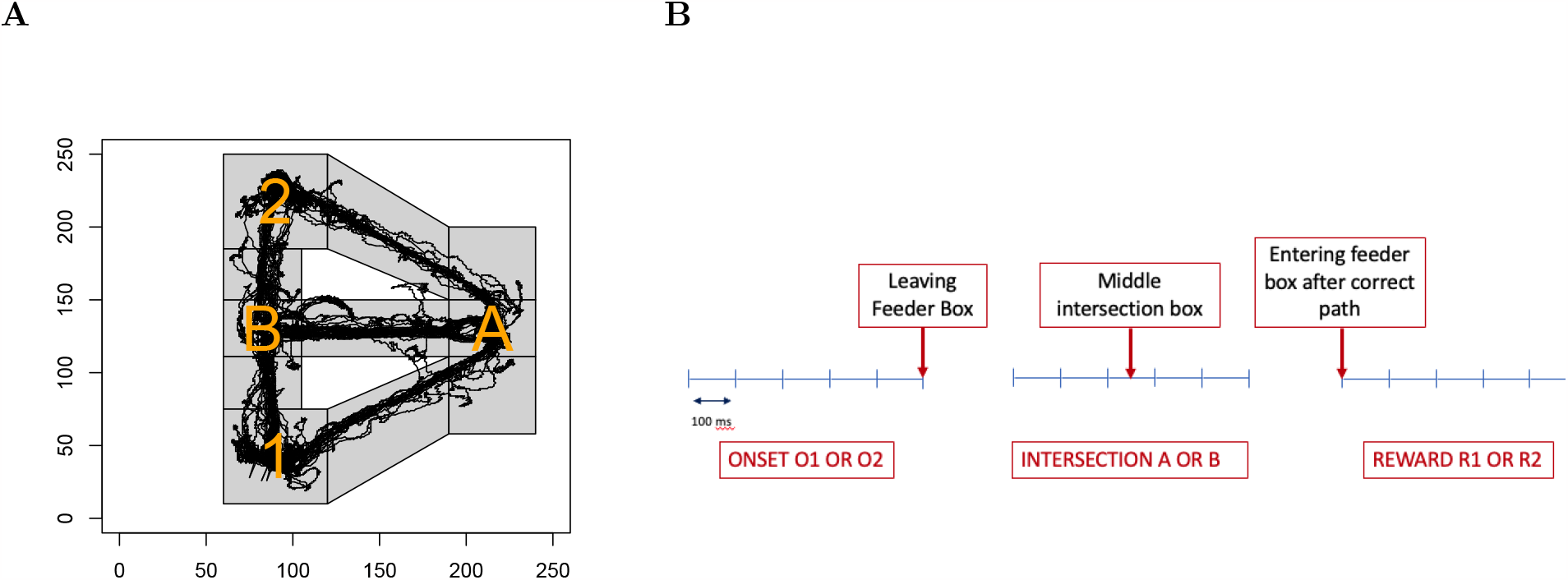
Timeline of a typical training session. **A**: The maze has been divided in boxes. The black line represents the trajectory of the animal during one example session. The feeders are located in the boxes 1 and 2. Maze intersections are located in boxes A and B. **B**: Each 500ms task event (centered on the onsets, intersections and reward locations) was divided in five 100ms bins. This leads to a total of 6 * 5 = 30 time bins.

**Figure 8:**
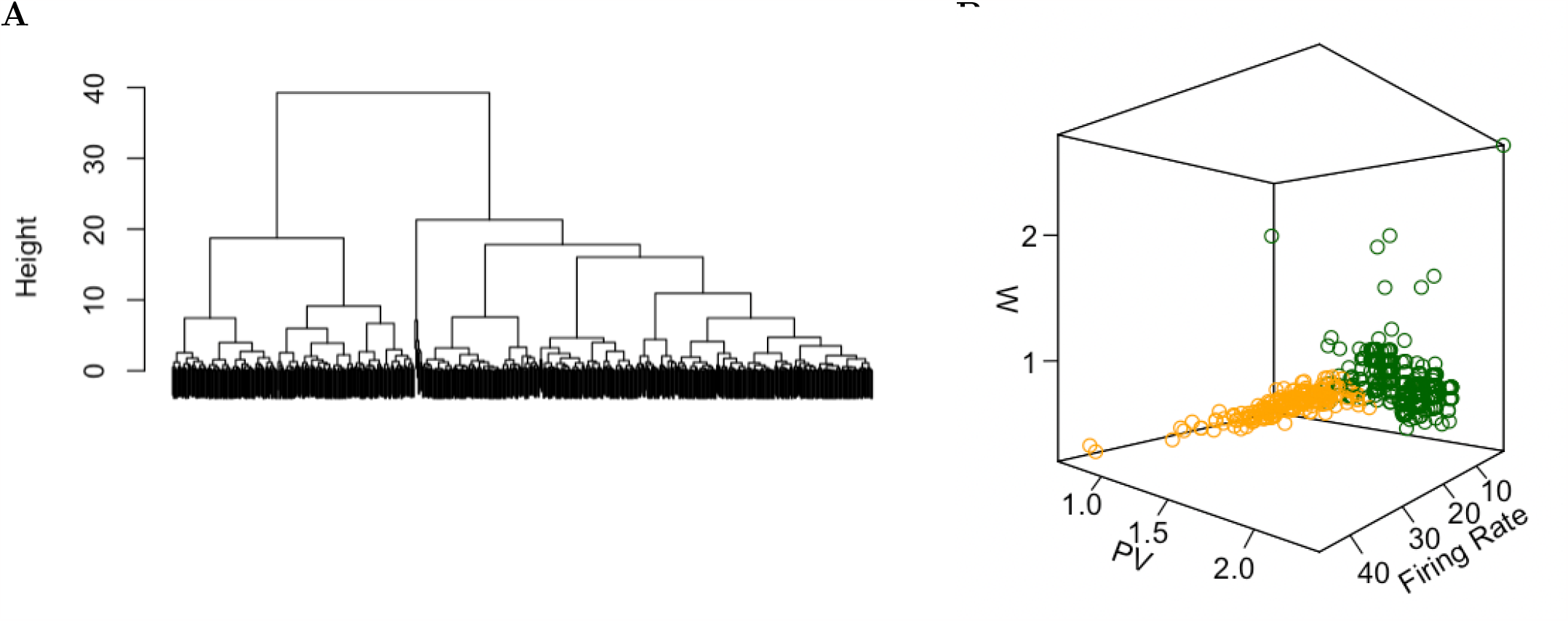
FSI-MSN classification. **A**: Dendogram of the hierarchical clustering (method “Ward D2”) of neurons based on waveform (wf) properties (Peak-Valley distance (PV), Width at mid-height (W)) and firing rate. Two clusters are identified using K-means method (with *K* = 2). **B**: 3D representation of wf properties and firing rate of the two clusters. Fast Spiking Interneurons (FSI) and Medium Spiny Neurons (MSN) are represented in orange and green, respectively. A total of 451 MSN and 199 FSI have been identified. No significant difference in the proportion of FSI/MSN was observed between DMS and DLS (p-value = 0.97).

